# Consensus or deadlock? Consequences of simple behavioral rules for coordination in group decisions

**DOI:** 10.1101/061010

**Authors:** Helen F. McCreery, Nikolaus Correll, Michael D. Breed, Samuel Flaxman

## Abstract

Coordinated collective behaviors often emerge from simple rules governing the interactions of individuals in groups. We model mechanisms of coordination among ants during cooperative transport, a challenging task that requires a consensus on travel direction. Decisions required for cooperative transport differ from other, well-studied consensus decisions because groups often deadlock, with individuals trying to move in opposing directions, and cooperative transport groups are often relatively small. Small groups may be more affected by individual nonconformity. Using deterministic and stochastic models, we investigate behavioral factors that affect deadlock duration. Our goal is to determine whether groups following simple behavioral rules can reach a consensus using minimal information. We define and investigate multiple types of behavioral rules that govern individual behavior and also differ in the information available. We find that if individuals more readily give up when they are going against the majority, groups rapidly break deadlocks. This occurs through positive and negative feedbacks that are implemented in our model via a single mechanism. We also find that to quickly reach a consensus, groups must have either a shared bias, high sensitivity to group behavior, or finely tuned persistence. While inspired by ants, our results are generalizable to other collective decisions with deadlocks, and demonstrate that groups of behaviorally simple individuals with no memory and extremely limited information can break symmetry and reach a consensus in a decision between two equal options

## Introduction

Workers of some ant species collaborate to carry objects many thousands of times their mass (1–5). For example, weaver ants, *Oecophylla longinoda,* carry dead vertebrates including birds and reptiles (2). Like many collective decisions, this requires a high degree of coordination across many individuals. The individual behavioral rules that lead to this coordination are difficult to identify empirically because they cannot be directly measured. Therefore, we use a proof-of-concept model (6) to examine the consequences of multiple behavioral rules individuals may use, and to see what individual characteristics lead to groups of effective transporters.

Across organizational scales, the patterns and complexity of many biological systems emerge from groups of individuals obeying relatively simple rules, often without a leader (7). The rules typically apply to individuals interacting with their neighbors, and exploit positive and negative feedback mechanisms leading to coordinated group dynamics. Of course rules do not have to be simple, but if robust, efficient coordination is possible with simple rules, then there is no reason for complex individual behaviors to evolve. Simple rules lead to group phenomena that are generally robust; when there is no leader, individuals are expendable, and the simplicity of the rules means that nearly any individual can perform appropriate behaviors with high fidelity and reliability.

Because of these benefits, interest in discovering rules for coordination has produced a rich literature, and there has been particular interest in group decision making (8–11). This includes nest-site selection decisions in honeybees and *Temnothorax* ants (12–15), decisions by groups of neurons in brains (16), decisions in non-neuronal organisms (17), and more. Ant colonies are particularly well suited to studies of collective behavior because workers can be easily observed and manipulated, and indeed, pheromone trail formation in ants is a classic study system for self-organized decision making (7).

One of the tasks groups of some ants pieces must complete is cooperative transport - the movement of large objects such as food items, intact, by multiple individuals (18). This is an interesting task for coordination research because it is prone to deadlocks, and has direct applications in robotics (4,19–22). A major challenge for cooperative transport is that the individuals must move the object the same direction. Ant species vary substantially in their ability to overcome this challenge. Some species move objects rapidly toward their nests, while many others are categorized as uncoordinated, with workers pulling in opposing directions for minutes or hours, making no progress (5,23). Prior research has revealed causes and effects of many aspects of cooperative transport, including selection pressures, ecology, recruitment, and more (reviewed in 4,5,18). This previous research has also included detailed descriptions and models of cooperative transport, and in some cases models have been compared with empirical data (e.g. 4,22,24). But these studies have not focused on comparing alternative behavioral rules for overcoming the coordination challenge; thus, our understanding of behavioral rules in cooperative transport is limited. Some investigators have suggested that ants in groups use the same rules as individual transporters (reviewed in 4). However, if rules for individual transport were sufficient, one would expect most ant species to be relatively efficient at cooperative transport. This is not the case, and it is reasonable to think that efficient cooperative transporters have behavioral rules tuned to this task. What behavioral mechanisms separate the coordinated from the uncoordinated transporters?

We use models to explore in detail the consequences of several broad categories of behavioral rules for coordination, including the information individuals must be capable of receiving to follow the rules. Our goal is not to identify the exact rules employed by all ants, but rather to explore the simplest behaviors and minimum information required for successful transport. Thus, we leave the comparison of our predictions with empirical patterns of transport for future research. Like other proof-of-concept models, the value in this work is that it tests the logic of verbal hypotheses and creates predictions that can be empirically tested (6). Our investigations generate hypotheses for cooperative transport adaptations and offer insights into consensus decisions in other groups. The broad modeling approach we employ has been used extensively to elucidate behavior that is difficult to measure in collective systems, including social insects, robots, and beyond (e.g. 25–30).

As discussed above, we do not yet understand what adaptations, including behaviors and sensory systems, allow some species to coordinate group transport efforts. However, we do know that cooperative transport efforts in uncoordinated species are characterized by many deadlocks, or periods in which individuals are attempting to move the object in opposing directions (5). Even in species with efficient cooperative transport, short-lived deadlocks occur, as groups first have a period of uncoordinated efforts before a consensus about travel direction emerges (4,5). Given that workers from the same colony are all ostensibly working to bring the object back to the nest, it is perhaps surprising that they so frequently pull in opposing directions. However, careful consideration reveals several naturally occurring circumstances that intuitively predict deadlocks. If workers arrive at the object from different nest entrances, each would likely attempt to move the object back to the entrance from which it emerged. We also expect deadlocks if multiple paths lead back to the nest or if individuals have conflicting information about the direction of the nest. Even during a cooperative transport effort that is already successful, a deadlock may occur if the group comes across an obstacle, as this requires a decision about which direction to move around the obstacle. Overcoming deadlocks, regardless of the cause, is crucial for cooperative transport.

Deadlocks are caused by workers being uncoordinated about travel direction (5,18). In a deadlock, some or all of the force that a worker imparts on the object is cancelled out by the forces of other workers. The resulting overall force is not large enough to allow for movement. Ants may also fail to move an object if an insufficient number of workers engage in transport, even if those workers are coordinated. We do not consider this situation to be a deadlock; in our usage, “deadlocks” occur only due to workers attempting to move the object in different directions. If group members can coordinate their forces, then they will rapidly break a deadlock and begin moving, while other deadlocks may last hours (23).

Cooperative transport is an especially interesting coordination task in part because of deadlocks. In other types of decisions, deadlocks may be unlikely or short-lived because early fluctuations are amplified by positive feedback (7). Also, large groups may be less likely to experience deadlocks, as they are less affected by single individuals. But cooperative transport groups can be as small as a few individuals, and deadlocks prevent transport in many species. Cooperative transport is also unusual because group members are physically tethered together by the object they are attempting to carry; split decisions are impossible. Groups must either break deadlocks or remain stuck. We evaluate the factors that affect the duration of deadlocks in cooperative transport, in a decision between two options – in our study, two travel directions. Specifically, we investigate the behavioral rules, information, and minimum complexity of individuals required in order to break deadlocks. Our approach consists of both deterministic and stochastic models of cooperative transport, in one spatial dimension. These models examine a decision between two options for direction of travel: left or right. Although real ants navigate in two dimensions, the one-dimensional problem is realistic in many scenarios, such as ants navigating along a defined pheromone trail or choosing which direction to move around an obstacle. We simulate transport under many conditions that vary in the behaviors of the individuals engaged in transport. Specifically, we model multiple broad categories of behavioral rules. These sets of rules differ in the kinds of information we allow individuals to perceive and the ways this information is used by individuals.

We use this approach to answer two primary questions. First, can realistic, simple behavioral rules reliably overcome deadlocks? As part of this question, we look at what information individuals must minimally receive. Second, what effects do persistence (maximum engagement time with the object) and sensitivity to information have on coordination? In answering these questions we generate hypotheses for cooperative transport adaptations and provide insight into the factors that affect deadlocks during cooperative transport, and during other collective decisions.

## Models

### Assumptions

We are interested in the minimum information and complexity requirements for deadlock breaking. We therefore assume individuals have minimal capabilities. As described below, we allow them only a few biologically plausible sources of information. Our simulated ants also have no memory, in that they do not use information from past experiences to shape future behaviors. We further simplify real cooperative transport efforts by assuming that all ants are identical.

Ants sense a wide range of stimuli (e.g. 31–33), though workers of a single species may be capable of sensing only a subset of the total possible information. There are several ways that workers in a cooperative transport group might gain information about what others in the group are doing. They could potentially communicate with one another, but while workers recruit additional help to the object to be carried, often with pheromone trails (18), there is no evidence of direct communication among ants after the recruitment phase. A more likely possibility is that workers communicate indirectly through the object being carried (4,19). Workers could sense the movements, deformations, and/or vibrations transmitted through the object, caused by the other workers' efforts (18,19,22). This indirect communication does not require an evolved signal, as workers are simply detecting physical cues that necessarily arise when forces are applied to an object. Hence, in terms of the kinds of information available in our model, we only consider cues and information transmitted through the object itself, including the size and direction of the overall force vector applied to the object. Our model thus serves as a logical test of this hypothesis regarding information transfer. We determine whether information easily transferred indirectly through the object is sufficient to bread deadlocks. Our assumptions are appropriate based on existing literature regarding complexity requirements for group decisions and hypotheses specific to cooperative transport (17,18).

#### Deterministic Model

We developed a deterministic, ordinary differential equations (ODE) model that simulates the average behavior of individuals. The model is Markovian – individuals have no memory – but non-linear. We model movement in one spatial dimension implicitly – we do not explicitly model the location of individuals or the object in space – and use continuous time and continuous abundances of individuals (but see individual-based model below). Individuals are identical, and the total number of individuals is fixed at 20; for some analyses we explored the effect of changing group size analytically and by evaluating groups with a total of 6 or 200 individuals. It is appropriate to have a fixed number of individuals because the number of workers that can participate in cooperative transport will be limited by the number of grasping points on the object. Furthermore, our assumed behavioral states allow varying numbers of individuals to be engaged with the object at any one time. Specifically, our model assumes that each individual occupies one of three mutually exclusive behavioral states: 1) trying to move the object to the left, 2) trying to move the object to the right, or 3) disengaged from the object (Fig 1). We do not distinguish between pushing and pulling; individuals pushing from the left and pulling from the right are both in the “move right” behavioral state. Individuals move from the disengaged state to an active state by “joining,” and from one of the active states to the disengaged state by “giving up.” These transition rates are important model parameters that govern the number of individuals (abundance) in each behavioral state over time. We look at these abundances to see the extent to which the group converges on a single direction under the parameters of a specific model run.

**Fig. 1:**
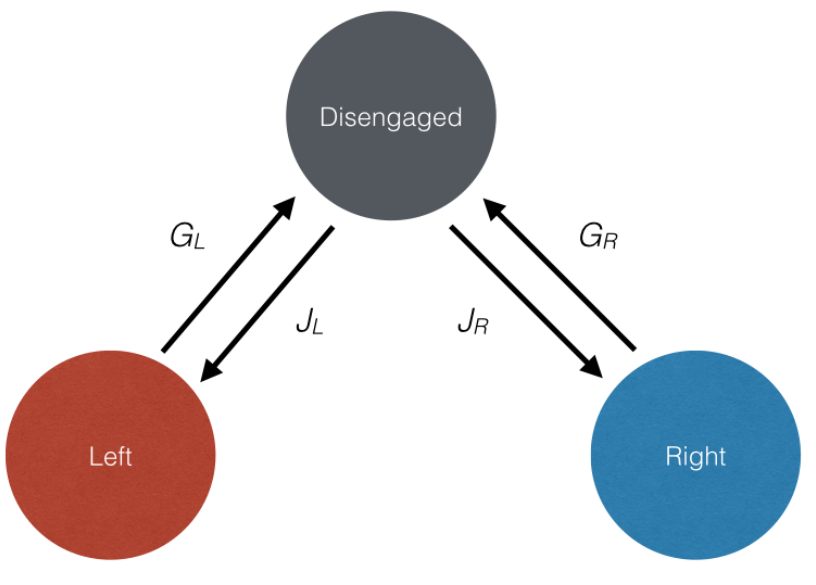
Model diagram. Individuals belong to one of three behavioral states: moving left, moving right, and disengaged. Individuals move between these states at rate constants *G_L_*, *G_R_*, *J_L_*, and *J_R_*

*Joining*. Disengaged individuals join the transport efforts to the left and right with rate constants *J_L_* and *J_R_*, respectively. The realized joining rates depend on the number of disengaged individuals; the instantaneous joining rate for the left state is *J_L_* multiplied by the number of disengaged ants. The joining rate constants do not change in time but may differ from each other, i.e. individuals may join the “move left” behavioral state at a higher rate than the “move right” state. If *J_L_* and *J_R_* are not equal, this ensures a directional bias, which is how we represent individuals having information about the direction of the goal.

In real ants, directional cues about the location of the nest come from one or more sources, such as pheromone trails, visual navigation, or path integration [5, 6]. Whatever the sensory modality may be, we assume this information is not perfect. That is, even if there is a directional bias, some individuals still choose the other direction (i.e., *J_L_, J_R_* > 0). Joining rate constants do not vary during the transport effort; for example, groups are not capable of altering their bias in favor of the “winning” direction (here we use the “winning” direction to indicate simply the direction that has more individuals). This makes sense given our conservative assumptions about individuals' memory and sensory capabilities: individuals that are disengaged, and therefore not in contact with the object to sense information transmitted through it, cannot perceive which direction is winning and have no memory about which direction was winning when they were last engaged.

*Giving-up*. Individuals in the active behavioral states (left and right) give up at rate constants *G_L_* and *G_R_*, respectively. We model three sets of behavioral rules for giving up rates. These sets of rules also differ in the kinds of information individuals act upon (Table 1). We do not suggest that all of the variation in cooperative transport behavior in ants is captured by these three sets of rules; rather, we explore these rules in an effort to see if such simple rules are sufficient to break deadlocks, and if so, under what conditions.

**Table 1:** Modeled sets of behavioral rules and information required for each.

|  | Rule(s) | Information used |
| --- | --- | --- |
| **Uninformed** | If in one of the active states, give up (become disengaged) with a constant rate | None |
| **Oblivious** | If in one of the active states, give up more readily when transport is unsuccessful, and less readily when it is “successful” (see text) | Must be capable of measuring the “success” (i.e. extent of coordination) but not the direction relative to ones own force input. |
| **Informed** | If in one of the active states, and if transport is successful, give up readily if going the opposite way as the majority and less readily if going the same way as the majority. | Must be capable of measuring (i) extent of coordination and (ii) preferred direction of the majority and must compare to the direction of one’s own force input. |

Behavioral rules differ among different model runs, but within one run of the model all individuals are identical and have the same rules and parameter values. In “uninformed” groups, giving up rate constants, *G_L_* and *G_R_,* are equal and do not change over the course of the transport effort. In “oblivious” and “informed” groups (defined in Table 1), realized giving up rates can change over the course of the transport effort based on the abundances of individuals in the two active behavioral states *(N_R_* and *N_L_)*. Giving up depends on the “success” of transport. “Success” is operationally defined here as a high extent of coordination, measured as the absolute value of *N_R_* - *N_L_* divided by the total number of individuals in the system. In other words, the extent of coordination is the degree to which individuals are *un*evenly distributed across the two active groups.

In oblivious groups individuals can measure success but they cannot detect if they are contributing to or detracting from that success. Individuals give up less frequently when |*N_L_* - *N_R_*| is high, i.e. when the magnitude of the overall force vector on the object is high, regardless of whether they are currently in the “right” or “left” state. Individuals are oblivious to their own contribution. If the transport is successful because many more individuals are trying to move the object to the left rather than the right, individuals moving right, who are going against the majority, still rarely give up. This would occur in ants if they were capable of measuring the magnitude of the overall force on the object (or a proxy), but not the direction; for example, by sensing movement speed.

In informed groups individuals are capable of detecting the same information as in the “oblivious” case, but additionally they can determine if their contribution is with or against the majority. Individuals give up less frequently when the force vector on the object is high only if the direction of that vector matches their own direction. For example, when *N_L_* – *N_R_* is strongly positive, individuals in the “move left” state give up infrequently while individuals in the “move right” state give up quickly. Equations governing the giving-up rate constants, *G_L_* or *G_R_*, under each set of rules are listed in Table 2 and examples of how these functions behave are illustrated in Figure 2. In addition to the variables *N_R_* and *N_l_*, functions for determining *G_L_* and *G_R_* depend on one or more parameters (Table 2). These parameters represent persistence and sensitivity, and are discussed below.

**Table 2:**
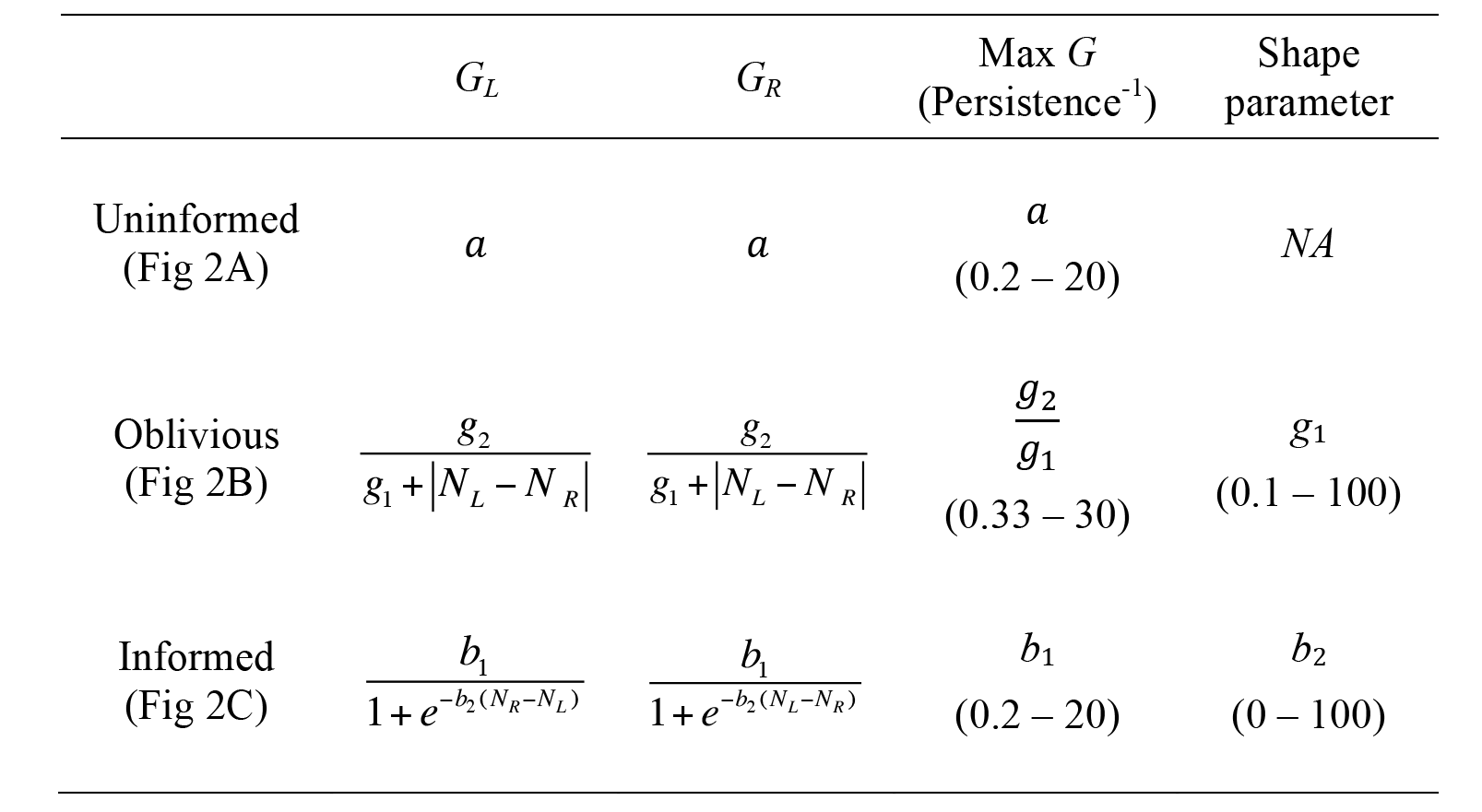
Functions governing the giving-up rate constants under each set of rules. Ranges of parameter values explored are in parentheses.

**Fig. 2:**
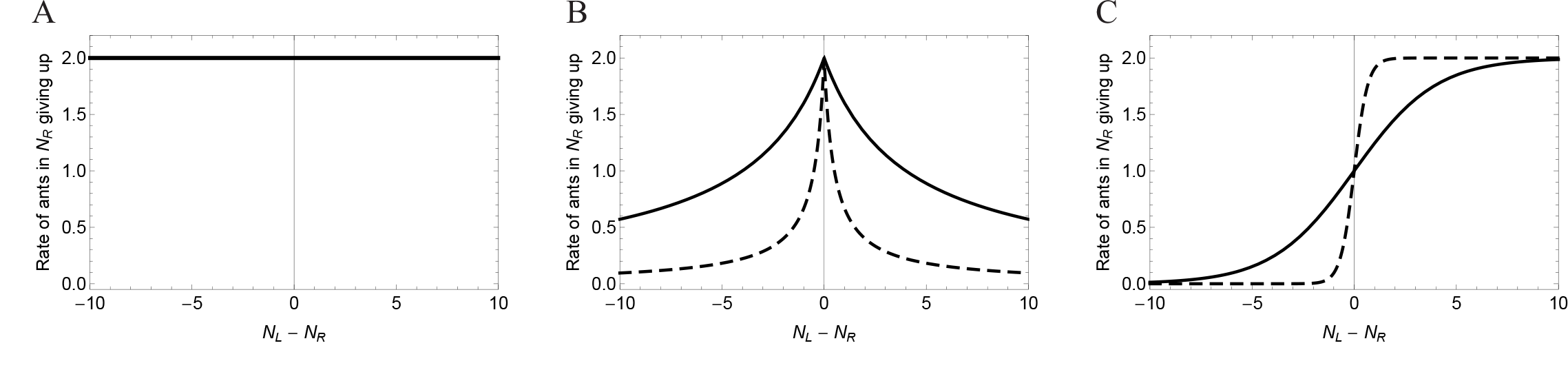
Giving-up rate constants for individuals in the “move right” behavioral state at various levels of success for each set of rules. The x-axis indicates a measure of success: the size difference between the two groups. (A) Uninformed rules, a = 2. (B) Oblivious rules, g_2_/g_1_ = 2, g_1_ = 4 (solid line) or 0.5 (dashed line). (C) Informed rules, *b_1_* = 2, *b_2_* = 0.5 (solid line) or 3 (dashed line). In (B) and (C), dashed lines indicate sharper shape parameters.

Note that *N_L_* – *NR,* is simply a measure of success that could be determined by ants in multiple ways. For example, large differences will correspond to larger speeds over ground, which an ant might measure by estimating optical flow, her own leg movements, or any other measure of speed. The direction of the force on the object, indicated by the sign of the difference between ants pulling to the left and to the right, could be estimated by comparing the direction of motion with the direction of an ant's own effort. Similarly, if the object is not yet moving, high values of *N_L_* – *N_R_* (or highly negative values) may correspond to deformations in the object, which ants could detect as their grasping point moving toward or away from them. The same quantities could potentially be estimated using other sensory modes.

*Persistence and sensitivity*. The giving up rates described above are tunable based on individuals' persistence and sensitivity to information. These parameters govern the shape and maximum values of the giving-up functions (Fig 2). This maximum giving-up rate is the inverse of the engagement time under conditions when individuals give up fastest (e.g., when *N_L_* = *N_R_* in the oblivious and informed cases). We refer to this engagement time as persistence.

Persistence is individuals' resistance to changing their behavior based on information (18), which could come from other individuals in the group, or other sources. Persistence can be measured in actual ants as the time it takes for an individual to give up or change the direction they are trying to move the object being carried. Highly persistent ants keep trying to move the object in the same direction for a long time, even without progress. On the other hand an ant with low persistence will try new strategies frequently, by pulling in different directions or even abandoning the effort temporarily or permanently. Intuitively, one expects a tradeoff for persistence. Groups with highpersistence may have long-lasting deadlocks, while groups may also deadlock if no individual is persistent enough. We look at the effect of persistence in our model by varying the maximum possible giving-up rate of the group. A highly persistent group still varies their giving-up rates under the “oblivious” and “informed” rules as described above, but for these groups the highest possible giving-up rate, which we impose, is low compared with groups with low persistence (Fig 2). We ran the model with each of many maximum giving-up rate constants to examine the effect of persistence on extent of coordination; higher maximum giving-up rate constants mean lower persistence and vice versa (Table 2).

For the oblivious and informed cases we can also tune the parameters to change the sensitivity of individuals to the success of transport, that is, the magnitude of |*N_L_* – *N_R_*|. We do this by changing the shape of the giving-up functions through manipulations of the shape parameters *(g_1_* and *b_2_*; Table 2), making the transition from low to high giving-up rate constant sharper or more gradual (Fig 2). With a gradual shape, small changes in success mean only small changes in the frequency of giving-up; individuals with a gradual shape therefore have low sensitivity to transport success. On the other hand, for sharp shapes, a small change in success may lead to a dramatic change in this frequency; this means individuals are highly sensitive. Differences in sensitivity could be caused by a number of factors, including error in sensing the group sizes. This shape parameter can be parameterized for real organisms by fitting functions to data on individuals, for whom cooperative transport efficiencies are experimentally manipulated.

*Differential equations*. The model consists of the following set of differential equations for the number of individuals in each behavioral state (moving left, moving right, disengaged):

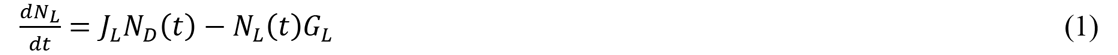

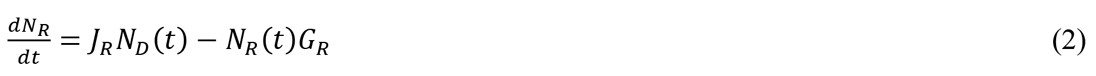

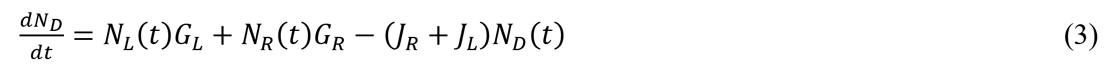

where *N_D_* is the number of individuals in the disengaged state (Fig 1). The ODE are nonlinear due to the dependence of *G_R_* and *G_L_* on *N_R_* and N_L_. There is a constant number of individuals total (i.e., *N_D_ + N_R_* + *N_L_* = *N* = constant), so this is a closed system, making the third differential equation implicit in the first two. Therefore in some cases we present results for the number of ants in the left and right states only. The ODE will always satisfy the following equations at equilibrium.

In the uninformed and oblivious cases, *G_L_ = G*_R_, so equation 6 simplifies to the following.

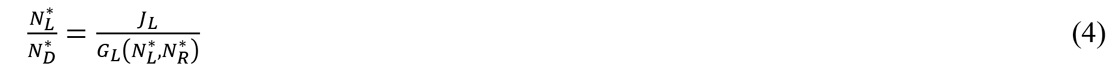

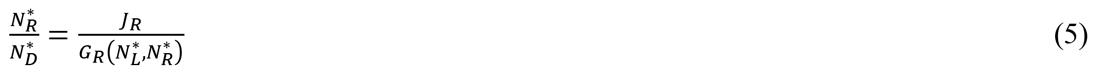

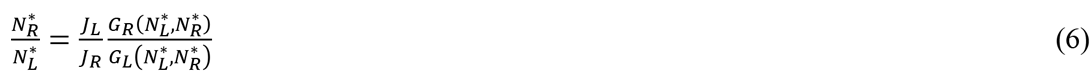

In the uninformed and oblivious cases, *G_L_ = G*_R_, so equation 6 simplifies to the following.

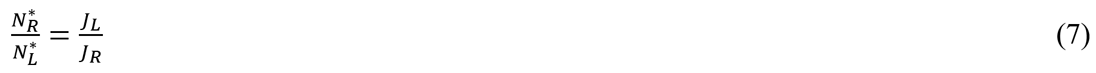

Because *G_R_* and *G_L_* are nonlinear functions of *N_R_* and *N_L_* in the oblivious and informed cases, it is difficult to solve this system differential of equations analytically. We numerically solved the ODE for each of nearly fifteen thousand sets of parameters, thus running the model under different sets of behavioral rules, global directional biases, and persistence and sensitivity. The range of parameter space explored for giving-up parameters is shown in Table 2. Additionally, we explored directional biases ranging from 0 to 0.89. We then queried the results for particular metrics of interest, including the maximum extent of coordination on a direction (unevenness in the distribution of individuals across the left and right groups). We obtained numerical solutions using *Mathematica* (version 9.0.1.0) and we analyzed our results using *Mathematica* and R (RStudio version 0.98.977). In addition to the numerical solutions, we analytically explored the stability of deadlocks in the informed case using fixed-point analysis (34, see S4 Appendix).

#### Stochastic Extension

Our ODE makes certain assumptions required for any ODE, including instantaneous updating of information and continuous, rather than discrete, individuals. To test whether our conclusions are robust to these assumptions, and to look at the potentially important influence of stochasticity, we extended the model to a stochastic framework. The stochastic extension is an individual-based model operating in discrete time. We converted the instantaneous joining and giving-up rate constants *(J_L_, J_R_, G_L_*, and *G_R_)* to probabilities of joining or giving up in a given time step with the equation

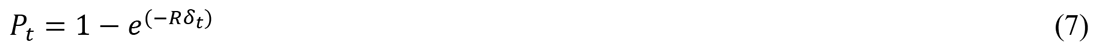

where *P_t_* is the probability of a behavioral shift in one time step, δ_*t*_ is the length of a time step (here, time steps were always unit length), and *R* is the instantaneous rate constant, either *J_L_, J_R_, G_L_*, or *G_R_*. We ran the stochastic simulation for 60 time steps; this duration was more than sufficient to capture transient dynamics. All other model assumptions and parameters were the same as in the deterministic model, including the three sets of rules.

In each time step we allow individuals to change their behavioral state. An active individual changes its status by giving-up with a probability equal to the giving-up probability for that individual's current state (left or right), and disengaged individuals can change their status by joining. Because disengaged individuals can change their status in one of two ways (joining the left group or the right group), we first calculated the joint probability of an individual joining at all. For individuals that were to join, we then stochastically determined whether they joined left or right using the relative probabilities of each. We ran the stochastic model under the same parameter sets as the deterministic model, querying 1,000 simulations for each set of parameters. As with the deterministic model, we examined the extent of coordination. We performed and analyzed stochastic simulations in R (RStudio version 0.98.977).

### Results

#### Deterministic Model

Our primary measurement of success is the extent of coordination, which is the difference in the number of individuals in the active behavioral states (left and right) divided by the total number of individuals in the system. If the transport is uncoordinated, there are roughly equal numbers of individuals pulling each direction, or most individuals are disengaged. Streamplot representations of the vector fields portray the dynamical behavior of the system in Figure 3. Panels in the figure show different parameter sets, corresponding to each set of behavioral rules with differing directional biases. The streamplots indicate the direction the system tends towards under all possible conditions for the numbers of individuals in each behavioral state *(N_L_* and *N_R_)*. The number of disengaged individuals, *N_d_*, is not shown explicitly because the total number of individuals is fixed at 20.

In the absence of a directional bias (*J_L_ = J_R_*), both the uninformed and oblivious rules have stable equilibria (Fig 3C, 3F). These are deadlocks, with equal numbers of individuals pulling left and right (as shown in equation 7). Because they are stable, perturbations away from these equilibria lead back to them (Fig 3C, 3F). In other words, with no directional bias the uninformed and oblivious rules have deadlocks that cannot be broken. In informed groups, however, the equilibrium is unstable even if *J_L_* = *J_R_* (Fig 3I). If a deadlock occurs in this case, small perturbations grow exponentially, leading to convergence on one direction, which breaks symmetry. Although an unstable equilibrium occurs across most of the parameter space for informed groups, with small values of the shape parameter *b_2_*, the equilibrium is stable and deadlocks are maintained. Thus there is a critical value of *b*_2_ at which a phase transition occurs, from stable to unstable equilibrium. Using fixed-point analysis (34) we analytically determined that this critical value occurs when *b*_2_ has the following value:

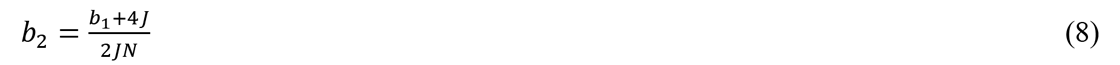

where J is the joining rate constant for each side (*J = J_L_ = J_R_*) and *N*is the total number of individuals in the system. This indicates that total group size affects deadlock breaking. Smaller groups require higher sensitivity (*b_2_*) to break deadlocks even in the informed case, and sensitivity is less important for large groups. Details of the fixed-point analysis are included in S4 Appendix.

**Fig. 3:**
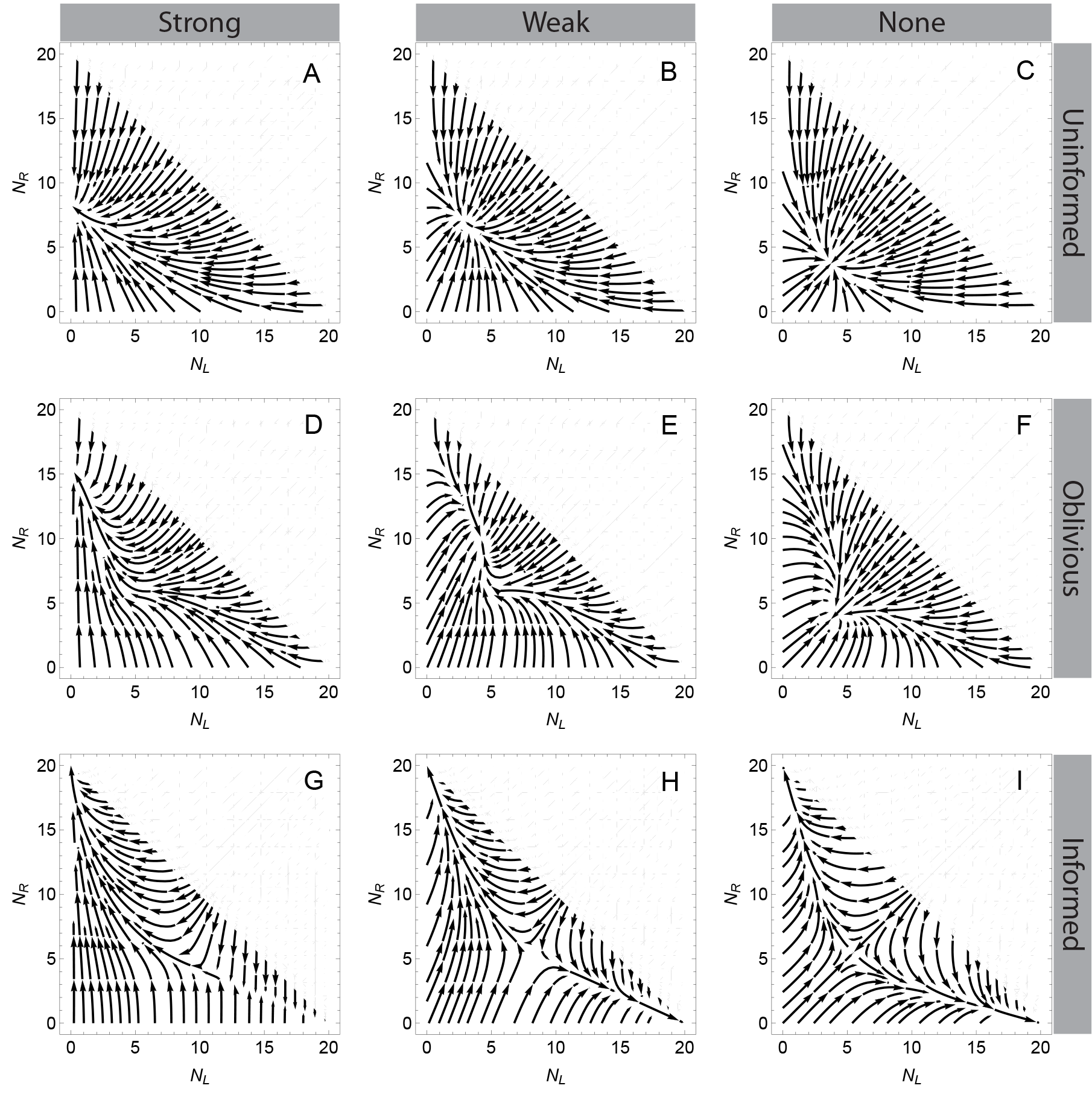
Streamplots of system dynamics, showing the direction the system tends towards for various abundances in each behavioral state. A-C: Uninformed rules, a = 1; D-F: Oblivious rules, *g*_1_ = 4, *g*_1_/*g*_2_ = 1; G-I: Informed rules, b_1_ = 1, b_2_ = 0.5. A, D, and G: Strong directional bias, *J_L_* = 0.01, *J_R_* = 0.7; B, E, and H: Weak directional bias, *J_L_* = 0.3, *J_R_* = 0.7; C, F, and I: No directional bias, *J_L_* = *J_R_* = 0.3.

When a directional bias is present *J_L_* ≠ *J_R_*) more individuals attempt to move the object in the direction favored by the bias, regardless of the set of rules (Fig 3, two left-most columns, also see equations 6 and 7). This is true regardless of the initial conditions for uninformed and oblivious groups; for informed groups, a large enough difference in the initial group sizes can overcome a joining bias (see rightmost portion of Fig 3G, 3H). The presence of a directional bias increases the extent of coordination, and, intuitively, strong biases lead to more coordination than weak ones (Fig 3 left column compared with middle column). However, for a given directional bias, individuals in informed groups are still more coordinated than individuals in uninformed or oblivious groups. Stable equilibria involving individuals working against one another still occur with a weak bias using these sets of rules (Fig 3B, 3E). With a sufficiently strong directional bias, in both uninformed and oblivious groups, the system moves to a state with almost no individuals going against the bias (Fig 3A, 3D), but there are still a substantial number of disengaged individuals who do not contribute to the effort (shown implicitly in Figure 3). This is because the disengaged group is constantly replenished by individuals giving up from the two active states. There are almost no disengaged individuals in informed groups. Thus, a directional bias allows for an unequal distribution of individuals between the two active states regardless of the behavioral rules, but the informed case still outperforms the other behavioral rules in that it maximizes engagement and the difference in group sizes.

#### Stochastic Model

The stochastic results match results from the deterministic model. Figure 4 shows the number of individuals in each behavioral state for two examples of stochastic simulations, under the same parameter sets shown in Figure 3. Histograms of the behavior across 1,000 simulations, at specific times, are shown in Figure 5 (also see S1 Movie). When a directional bias is present more individuals try to move the object in that direction than in the other direction under our initial conditions of all individuals beginning as disengaged. In the absence of a directional bias, roughly equal numbers of individuals are in each active state in uninformed and oblivious groups, while individuals converge on either direction in informed groups. In each of 1,000 simulations, the informed case allowed for convergence to a pure state (every individual or nearly every individual in the system pulling the same direction) even with no directional bias (Fig 5 and S1 Movie). On the other hand, oblivious groups perform no better than uninformed groups, and neither of these sets of rules ever allowed for convergence on one direction.

**Fig. 4:**
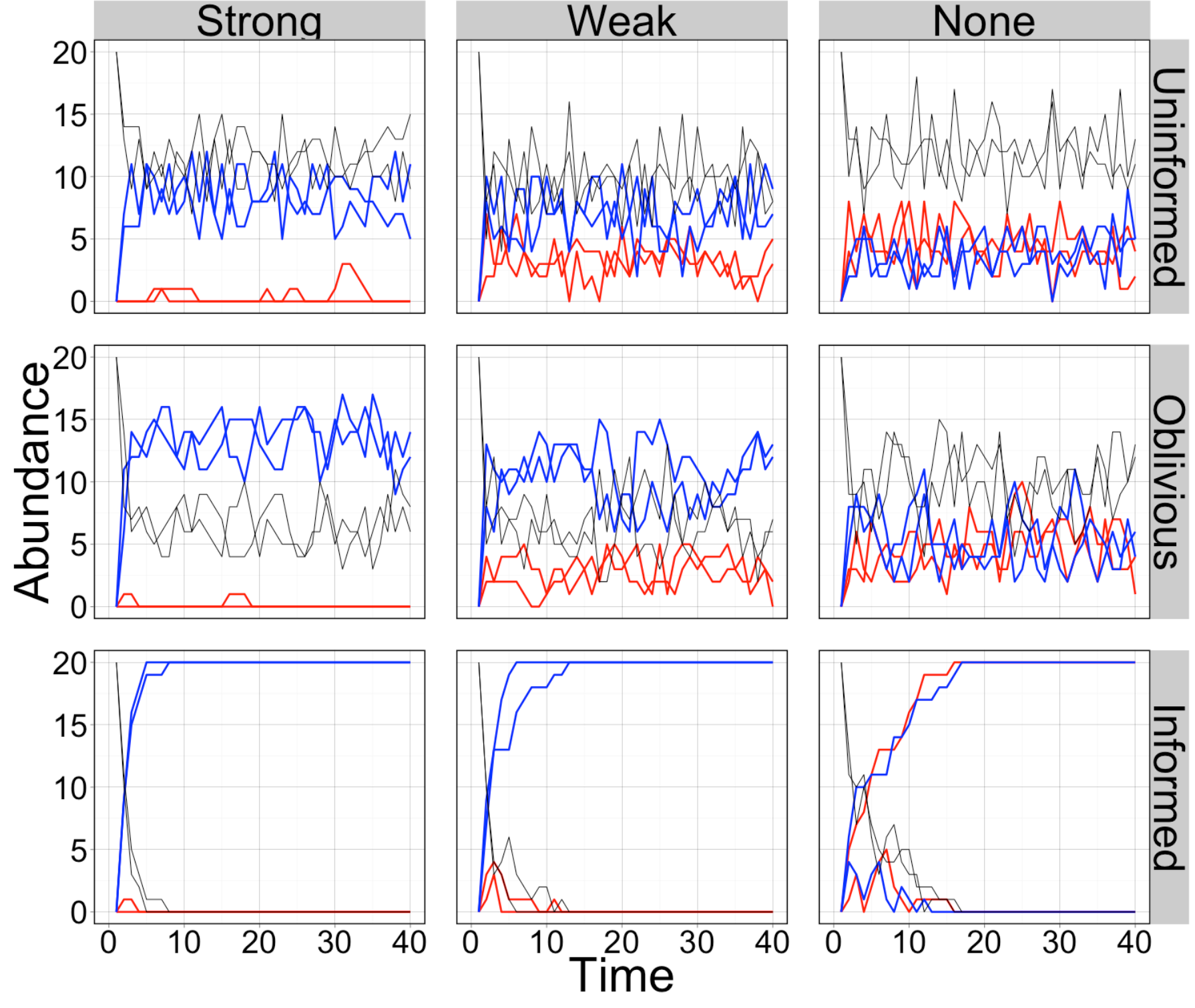
Abundance of ants in each behavioral state over time, for two example simulations with each set of parameters. Blue: moving right, Red: moving left, Black: disengaged. Columns are different directional biases and rows are different sets of behavioral rules. The parameter values are the same as the analogous panels in Figure 3. Uninformed rules: a = 1; oblivious rules: *g*_1_ = 4, g_1_/g_2_ = 1; informed rules: b_1_ = 1, b_2_ = 0.5. Strong directional bias: *J_L_* = 0.01, *J_R_* = 0.7; weak directional bias: *J_L_* = 0.3, *J_R_* = 0.7; no directional bias: *J_L_* = *J_R_* = 0.3.

**Fig. 5:**
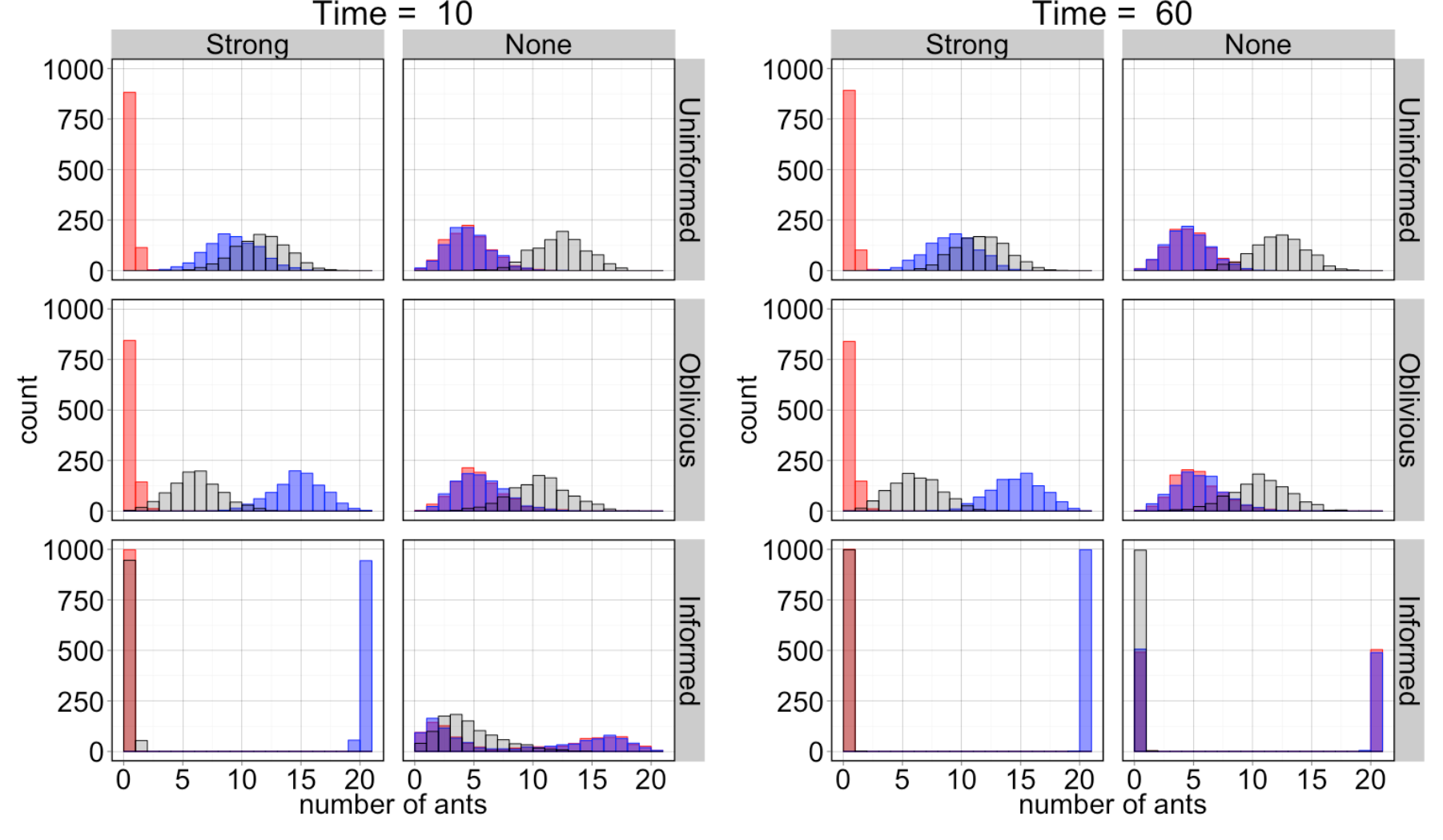
Histograms showing the state of 1,000 simulations at given time points. The x-axis shows the number of ants, and the y-axis shows the number of simulations for which the given behavioral state had that many ants at that time. Blue bars are for ants moving right, red bars are for ants moving left, and black bars are for disengaged ants. Bars appear purple when red and blue overlap. The parameter values are the same as the analogous panels in Figures 3 and 4. Uninformed rules: a = 1; oblivious rules: g_1_ = 4, g_1_/g_2_ = 1; informed rules: b_1_ = 1, b_2_ = 0.5. Strong directional bias: *J_L_* = 0.01, *J_R_* = 0.7; no directional bias: *J_L_* = *J_R_* = 0.3.

When a directional bias is present, the informed case still leads to strikingly different performance than either of the other sets of rules. Individuals converge rapidly in informed groups, while in oblivious or uninformed groups convergence, which we define as an increasing coordination through time until all individuals are pulling the same direction, does not occur. There are more individuals pulling in the direction of bias but conditions do not improve over time (Figs. 4, 5, and S1 Movie).

#### Effect of persistence and sensitivity

Figure 6 shows the effect of persistence – or maximum engagement time – on the extent of coordination in the deterministic model for groups with total size fixed at 20 (see S2 Figure for results for other group sizes). The extent of coordination reported is the maximum observed over the time period evaluated. Parameter sets that converge more quickly on a direction will have a higher extent of coordination in that time period, and shorter deadlocks. Results in Figure 6 are therefore comparable across parameter sets, with higher agreement indicating more efficient transport. In the deterministic case the small perturbations away from equilibrium that lead to convergence in informed groups do not occur, so with no directional bias there is no coordination.

**Fig. 6:**
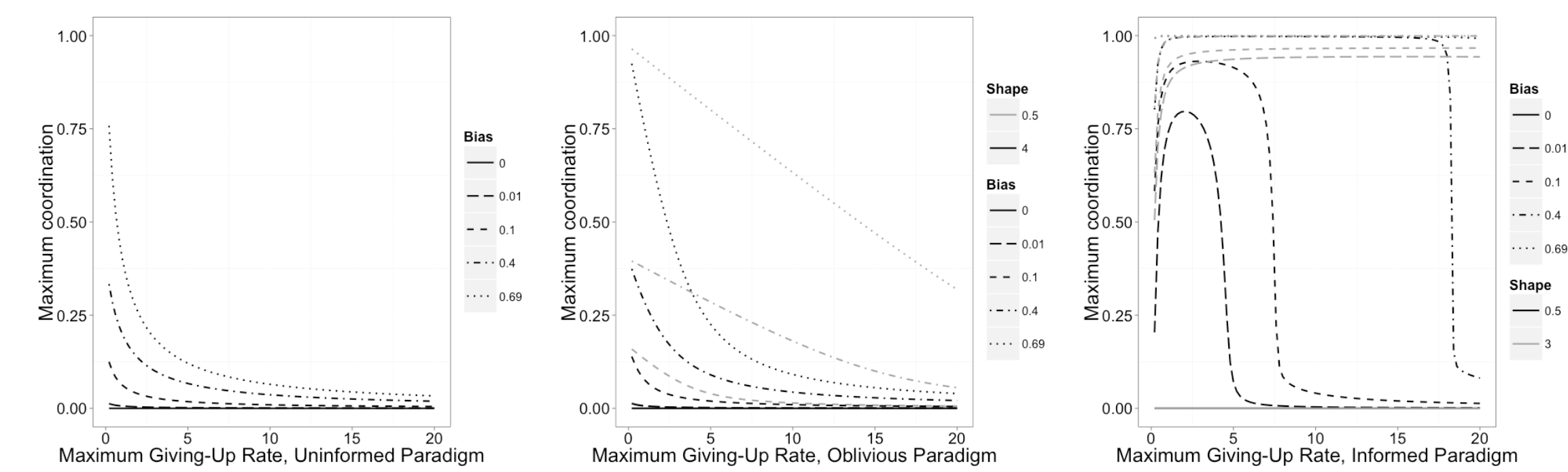
Effect of persistence (inverse of maximum giving-up rate constant) on maximum coordination. Maximum giving-up rate constant is the maximum possible as defined by the function, actual values will depend on the number of individuals in each group. Extent of coordination is defined as the difference in the number of individuals pulling right and left, divided by the total number in the system. Maximum coordination is the maximum observed over a given time period, rather than an absolute maximum; higher values on the y-axis indicate faster convergence. (A) uninformed rules, (B) oblivious rules, (C) informed rules. Lines with smaller dashes indicate lower directional bias. Gray and black lines indicate “sharp” and “gradual” shapes (sensitivities), respectively. Parameter values for shape match those in Fig 2.

The effect of persistence depends on the behavioral rules (Fig 6). In uninformed and oblivious groups, being highly persistent – having a low maximum giving-up rate constant – increases coordination (Fig 6A, 6B). In informed groups there is an optimal persistence value that maximizes coordination. The extent to which persistence affects coordination is stronger for small directional biases; at high directional biases there is a wide range of persistence values that result in high coordination. These results were not qualitatively different for different total group sizes, except that sensitivity, or sharpness of the giving-up function, was more and less important for smaller and larger groups, respectively (S2 Figure).

For oblivious and informed groups, the sensitivity changes the effect of persistence (Fig 6C and Fig 7); the uniformed case has no sensitivity parameter. In the oblivious case, sharper functions (lower values of *g*_1_) increase coordination for a given persistence value. In the informed case there is a critical sensitivity below which deadlocks cannot be broken, as discussed above and in S4 Appendix. This threshold depends on group size. Above this threshold, sharper functions (higher values of *b*_2_) further increase coordination, which has the effect of widening the range of persistence values that lead to coordination. For a moderate group size of 20 individuals, with a gradual shape and a small directional bias, there is a narrow range of persistence values that allow for high coordination. At small group sizes only groups with higher sensitivity or relatively strong directional bias coordinate successfully regardless of persistence, while large groups successfully coordinate across a wide range of persistence regardless of sensitivity and bias (S2 Figure). Figure 7 shows in detail the extent of coordination for moderately sized groups with a wide range of directional biases and persistence values for two shape values, both relatively gradual (see S3 Figure for small and large groups).

**Fig. 7:**
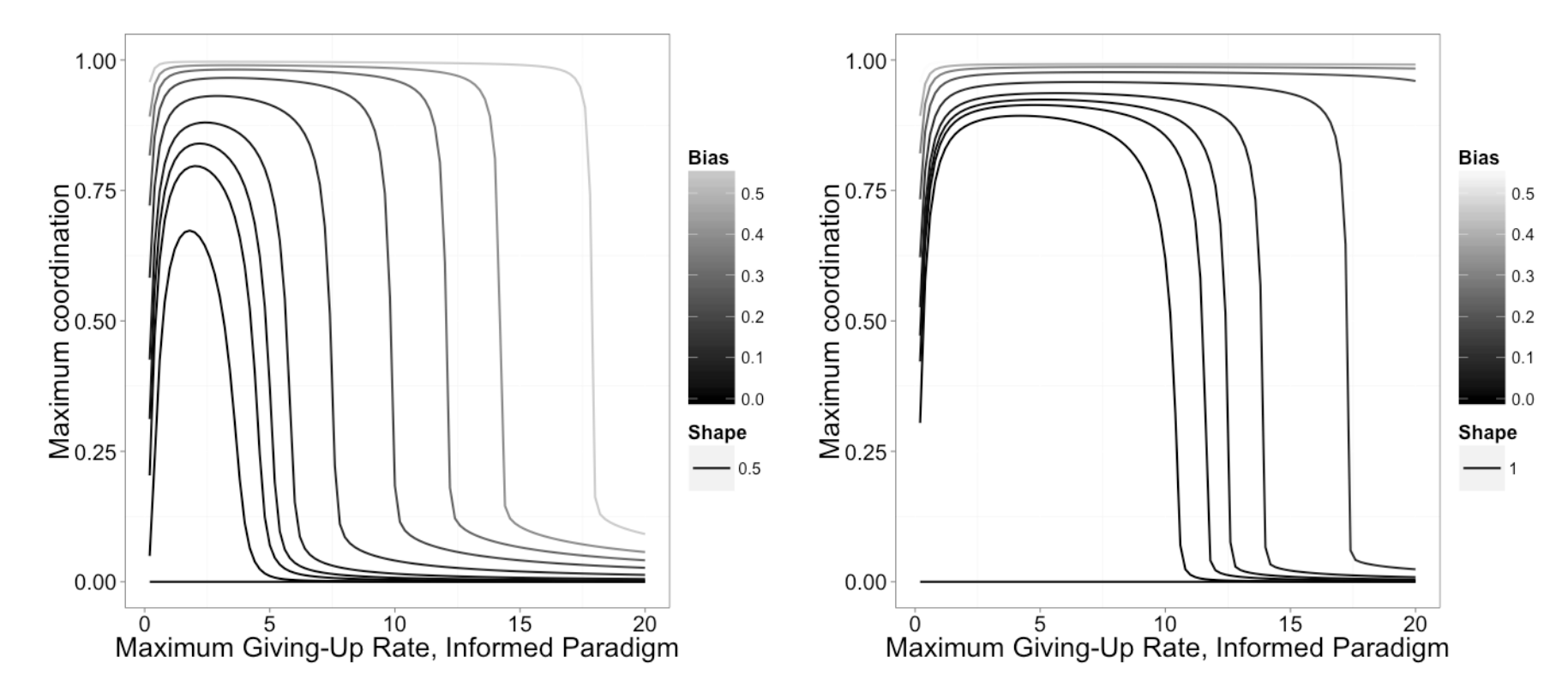
Effect of persistence (inverse of maximum giving-up rate constant) on maximum coordination in informed groups at low (gradual) shape values. Maximum giving-up rate constant is the maximum possible as defined by the function, actual values will depend on the number of individuals in each group. Extent of coordination is defined as the difference in the number of individuals pulling right and left, divided by the total number in the system. Maximum coordination is the maximum observed over a given time period, rather than an absolute maximum; higher values on the y-axis indicate faster convergence. (A) shape parameter, *b*_2_ = 0.5, which corresponds to the solid line in Fig. 2C. (B) *b*_2_ = 1, which is less gradual.

### Discussion

Can relatively simple individuals with minimal information break deadlocks? Our results show that, indeed, individuals with simple behavioral rules and no memory can break deadlocks. However, only individuals in our informed case convincingly succeeded. These individuals followed simple rules: 1) give up more readily if one is moving against the majority and 2) do this to a greater extent for extreme majorities than slight majorities. Using these simple rules, with minimal information available, groups rapidly converge on a single travel direction. Our deterministic and stochastic models agree, despite being formulated differently and having contrasting assumptions about individuals and time. This suggests that our conclusions are robust to specifics of model formulation.

In terms of information, it is sufficient for coordination for individuals to only be capable of measuring the direction that the majority of the group is trying to move the object and the relative sizes of the groups moving each direction (or a proxy, such as speed). This information is crucial; with insufficient sensitivity to these group sizes (low *b*_2_) groups do not form a consensus. The magnitude of the overall force on the object is a proxy of the relative group sizes (if the sizes of the two groups are approximately equal, the overall force vector will be small, as most of the force of one group is cancelled out by the other), and there are other possible proxies for relative group sizes. Ants may gather information about forces or group sizes by sensing the motion of the object itself, or if the object is not moving, by sensing vibrations or deformations in the object. If this is the case, a single sensory mode may provide all necessary information in informed groups. In nature, ants may have other information available, or may use different behavioral rules, but we show that by using these simple rules, groups are successful. Behavioral complexity comes at a cost, in terms of energy and information, so given that a simple solution exists it is likely that real ants adapted for cooperative transport also use simple rules.

If individuals have global directional cues that correspond to a shared directional bias, this helps promote coordination regardless of the other information available. Additionally, if there is only one correct direction, for instance if there is a single nest entrance, a shared bias towards the nest would help ensure the group converges on the appropriate direction. But directional bias is neither necessary, nor sufficient, for convergence on a decision.

These results make sense considering the high variation in cooperative transport ability among ant species. We expect workers of all species to be good at knowing the direction of the nest. So we expect directional biases to be common among species, at least for situations with only one correct direction. Considering that efficient cooperative transport is comparatively rare among ants (5,35), the presence or absence of directional biases is not a good explanation for the observed variation in efficiency. On the other hand, the behavioral rule of giving up more readily when an individual is moving against the majority is a potential adaptation that dramatically improves efficiency. Future research should test whether efficient transporters have this adaptation.

Our second question asked what effects do persistence and sensitivity (the sharpness of the giving-up function) have on coordination? These effects are complex and depend on the total group size and the behavioral rules. In the uninformed and oblivious cases, groups are most coordinated if individuals are highly persistent. While somewhat surprising, this makes sense in light of a tradeoff in persistence. Groups of highly persistent individuals may pull in opposing directions for a long time, but if movement does occur, either because of a directional bias or due to random fluctuations, the progress continues; they are unlikely to change their direction.

This suggests that high persistence allows species without other adaptations for cooperative transport, for instance those with behavioral rules similar to our uninformed or oblivious rules, to at least sometimes succeed at bringing a large object home to the nest. In such species, individuals are equally likely to give up whether they are helping or hurting the effort; even when successful movement occurs, individuals pulling with the motion may give up. High persistence makes it less likely that anyone will give up, allowing existing movement to continue. If, as in our model, individuals are identical, the individuals going the wrong way will also be unlikely to give up, so to minimize the length of deadlocks there should only be a small number of these individuals. A sufficient directional bias would accomplish this, and directional biases should be common in many circumstances (such as if the object is relatively far from the nest). So if high persistence is paired with a directional bias, it may allow ant species with rudimentary behavioral rules to conduct cooperative transport. Analogously, agents involved in any decision between two options, when they are unable to determine which option is winning, should be persistent to maximize the chance that a single option will be chosen.

In contrast to these results, in the informed case there is an optimum persistence value; groups with individuals more or less persistent than this value will be less coordinated. But the importance of persistence depends on directional bias, on the sharpness of the giving-up function, and on the group size. In most of the parameter space of our model, the range of persistence values that lead to high coordination is wide. Only when the directional bias is low and the sensitivity is above the critical threshold but still gradual does one find a narrow peak in coordination around the optimum persistence. This was especially true for smaller group sizes. Large groups had a wide range of persistence values that would lead to coordination regardless of sensitivity, indicating that it may be easier to coordinate in a large group rather than a small group. This makes sense given that small groups will be more affected by the behavior of single individuals. In order for informed individuals in groups of small to moderate size to be highly coordinated, they must have one, but do not need more than one, of the following: high directional bias (one option clearly favored over the other), high sensitivity to the sizes of the two groups, or finely-tuned persistence. Each of these is a potential adaptation for efficient cooperative transport in informed groups. This flexibility makes the behavioral rules in the informed case relatively robust to deficiencies in the individuals' capabilities as long as they have at least minimal accuracy in sensing group sizes.

The phase transition we observed in the shape parameter for the informed case indicates that individuals must have some threshold level of sensitivity of the sizes of the groups in order to reach a consensus. With more gradual shapes, the giving-up function approaches linear, and the less sensitive individuals are to the difference in size of the two groups. The phase transition makes intuitive sense, as the giving-up function becomes more linear, individuals switch too frequently to overcome deadlocks.

Because we did not constrain our model by tuning it to a particular species, our results are applicable to other collective decisions, especially those that are subject to deadlocks. A system in which groups must decide among multiple options is vulnerable to deadlocks, especially when the options are relatively equal (analogous to having no directional bias); small group size may also make deadlocks more likely. One of the best studied examples of collective decisions is nest-site selection in social insects (reviewed in 12). In fact, some recent work on the “stop-signal” in honeybees focuses on how this signal prevents deadlocks in nest-site selection (15,30,37).

The outcome of our deterministic model with respect to the effect of behavioral rules looks similar to the results of Seeley et al. (15) and Pais et al. (30), who each investigated decision-making dynamics in honeybee nest site selection with similar models (for example, compare Fig 3C to Fig 3 in 35 and inset in 27 Fig 2). Both models investigate the accumulation of “votes” for one of two, mutually exclusive choices in a decision, and in each case the number of individuals aligned with the two options determines which option is chosen. A key difference between the models, however, involves the timing of the decision. In honeybee nest-site selection, a decision is reached when a quorum of scouts is present at one of the potential nests (13). In cooperative transport, a decision is reached when the difference between the number of individuals in each group reaches a certain threshold – enough to begin movement – rather than when the absolute number of individuals in a particular group is high. Another key difference is the consequence of error. In honeybee nest site selection, the stakes are high. It is possible for a quorum to be reached at multiple potential nest sites at once, which may lead to the colony splitting, dramatically reducing the chance of survival of the colony (13,38). Because of the fact that the *difference* between group sizes is what matters in cooperative transport, such a split-decision is impossible.

But perhaps the most important difference between these models relates to communication. Unlike our model the Seeley et al. (35) and Pais et al. (27) models include direct communication among individuals. Honeybee scouts actively advertise for a particular nest site (a positive feedback mechanism) and stop other scouts from advertising for a different site using the stop signal (a negative feedback mechanism; 35). Our model produces some similar dynamics without direct communication through evolved signals; instead, individuals pick up on group cues. In informed groups, positive and negative feedback mechanisms are combined into a single mechanism that requires no signals. An individual is less likely to give up if her faction is large compared to the other faction (positive feedback), and more likely to give up if the opposite is true (negative feedback, analogous to cross-inhibition). Informed individuals only need to measure the relative group sizes to make effective decisions.

The Seeley et al. (35) and Pais et al. (27) models elegantly and realistically reproduce the dynamics of nest-site selection in honeybees. Our model is simpler, yet produces similar dynamics in terms of the accumulation of votes for a single option, indicating that communication among individuals is not necessary for a decision in the case of cooperative transport. The fact that some of our results are similar lends credence to the idea that results from one collective decision-making system can be generalizable to others. Among collective systems, social insects are uncommonly apt for experiment, since individuals are easily observed and manipulated. Because lessons are transferable across at least some systems, we can use social insects as model systems for other systems that are harder to study, like neuronal networks and immune systems.

Our model demonstrates that simple behavioral rules can lead to a consensus about travel direction during cooperative transport, even without a directional bias. Our simulated ants had no memory, limited sensory ability, and followed only simple rules, yet made decisions rapidly in informed groups. We identify a potential adaptation – giving up more readily when going against the majority – that allows for deadlock-breaking, and may explain why we see such large variation in cooperative transport ability among ant species. While it is currently not possible to directly measure this adaptation in ants, the consequences we have modeled here can, and should, be measured to see if real ants use this behavioral rule. Our model reproduces dynamics similar to those of other decision-making processes (15,30), and our conclusions are generalizable to other collective decisions, especially those prone to deadlock. Though cooperative transport is a challenging task that requires coordination, behavioral complexity is not a prerequisite for success.

## Acknowledgements

We thank James Marshall and Andrew Koller for advice on the model and implications. Ted Pavlic provided extensive and valuable comments on the model and manuscript. We also thank Tim Szewczyk, the Modeling Group, QDT, the Writing Cooperative, and the Breed Lab at the University of Colorado for comments on the model, manuscript, and code.

